# Viral-mediated expression of Erk2 in the nucleus accumbens regulates responses to rewarding and aversive stimuli

**DOI:** 10.1101/2023.10.03.560689

**Authors:** Lyonna F. Parise, Sergio D. Iñiguez, Brandon L. Warren, Eric M. Parise, Ryan K. Bachtell, David Dietz, Eric J. Nestler, Carlos A. Bolaños-Guzmán

## Abstract

Second-messenger signaling within the mesolimbic reward circuit is involved in both the long-lived effects of stress and in the underlying mechanisms that promote drug abuse liability. To determine the direct role of kinase signaling within the nucleus accumbens, specifically mitogen-activated protein kinase 1 (ERK2), in mood- and drug-related behavior, we used a herpes-simplex virus to up- or down-regulate ERK2 in adult male rats. We then exposed rats to a battery of behavioral tasks including the elevated plus-maze, open field test, forced-swim test, conditioned place preference, and finally cocaine self-administration. Herein, we show that viral overexpression or knockdown of ERK2 in the nucleus accumbens induces distinct behavioral phenotypes. Specifically, over expression of ERK2 facilitated depression- and anxiety-like behavior while also increasing sensitivity to cocaine. Conversely, down-regulation of ERK2 attenuated behavioral deficits, while blunting sensitivity to cocaine. Taken together, these data implicate ERK2 signaling, within the nucleus accumbens, in the regulation of affective behaviors and modulating sensitivity to the rewarding properties of cocaine.

## 1. Introduction

The mesocorticolimbic dopamine system, comprised of the ventral tegmental area (VTA), nucleus accumbens (NAc), and prefrontal cortex (PFC), is best known for its role in mediating the rewarding and reinforcing properties of drugs of abuse (Chao and Nestler, 2004; Holly and Miczek, 2015; Koob, and Kreek, 2007) however, given its prominent role in stress-responding, dysregulation of this circuit promotes aberrant mood-related behavior (Russo and Nestler, 2013). Indeed, clinical evidence also supports that perturbations in NAc function are present in individuals chronically suffering from affective (Ding 2022) and substance use disorders (Engeli, 2021; Wang, 2019). Interestingly, dopamine release from the VTA to the NAc has been shown to occur in response to both rewarding and aversive stimuli (Carlezon and Thomas, 2008; Nestler and Carlezon, 2006) making this a highly complex circuit capable of mediating maladaptive behaviors. Major depressive disorder (MDD) and drug use often co-occur in a bi-directional manner such that drug abuse increases the likelihood of developing mood disorders, and conversely, individuals with MDD are at an increased risk of developing drug-seeking tendencies (Burke and Miczek, 2013; Miczek et al., 2008). This suggests that there are overlapping stress-induced mechanisms that promote behavioral alterations and facilitate drug reward (i.e., cross-sensitization). Indeed, this perspective is supported by rodent models of stress-induced mood disorders and drug taking (Berton et al., 2006; Krishnan et al., 2007; Markou et al., 1998; Naranjo et al., 2001; Nestler and Carlezon., 2006). For example, both a priming injection of cocaine and exposure to an acute stressor are capable of reinstating drug-related behaviors in a rodent trained to extinguish drug seeking (Buczek et al., 1999; Erb et al., 1996). Furthermore, chronic stress appears to induce enduring changes in molecular and cellular processes that resemble those observed after extended exposure to various drugs of abuse (Bruijnzeel et al., 2004; Carlezon et al., 2005; Shaham et al., 2000; Willner, 2017).

While the precise mechanism(s) underlying these similarities remains elusive, second messenger signaling pathways have been repeatedly implicated in mediating at least some of these processes (Alcantara et al., 2014) (Altar, 1999; Berhow et al., 1996). Of particular interest is the mitogen-activated protein kinase, extracellular-regulated protein kinase (ERK), which consists of two isoforms, ERK1 and ERK2. It is hypothesized that these two isoforms have differing functions in the brain, however their discrete roles have not been fully characterized. The limited differentiation of their roles in studies can be attributed to the significant genetic sequence overlap between the two isoforms and the current inability of pharmacological tools to distinguish between ERK1 and ERK2 (Nishimoto and Nishida, 2006; Selcher et al., 2001). Nevertheless, several studies have suggested that in comparison to ERK1, ERK2 is more directly involved in drug-related mechanisms (Berhow et al., 1996; Einat et al., 2003; Iñiguez et al., 2010; Lu et al., 2005). Additionally, ERK2 has been implicated in the adaptive response to stress and is considered an integral mediator of effective antidepressant treatment, further supporting its role in modulating affective behaviors (Gourley et al., 2008; Iñiguez et al., 2010; Qi et al., 2008; Warren et al., 2011).

Although the function of ERK2 signaling within the VTA and other brain areas is relatively well understood (Berhow et al., 1996; Iñiguez et al., 2010; Warren et al., 2011), less is known about the role of ERK2 signaling within the NAc in the context of drug-related behaviors. Investigating this is important, as there is substantial evidence suggesting that intact NAc function is necessary for appropriately encoding the valence of drugs of abuse which can in turn influence drug-seeking behaviors (Namburi et al., 2016). Moreover, it’s worth noting that both stress and drug exposure has been shown to alter the morphology of NAc medium spiny neurons (MSNs), thereby influencing overall mesocorticolimbic communication, making this a particularly important phenomena to functionally assess in relation to kinase signaling. The goal of the present study was to determine the effect of directly manipulating ERK2 activity within the NAc on behavioral measures of mood and sensitivity to drugs of abuse, namely cocaine, while also exploring the effect of ERK2 modulation on the morphology of MSNs.

## 2. Methods

### 2.1 Animals

Male Sprague Dawley rats weighing 275–300 g upon arrival (Charles River Laboratories) were used in this study. Rats were housed in clear polypropylene boxes containing wood shavings in an animal colony maintained at 23–25°C, on a 12 h light/dark cycle in which lights were on between 7:00 A.M. and 7:00 P.M. Food and water were provided *ad libitum*. All procedures were in strict accordance with the *Guidelines for the Care and Use of Mammals in Neuroscience and Behavioral Research* (National Research Council, 2011) and approved by the Mount Sinai, University of Colorado-Boulder, and Texas A&M University Animal Care and Use Committees.

### 2.2 Chronic Unpredictable Stress

Chronic unpredictable stress (CUS) is a 4-week paradigm consisting of exposing rats to one stressor/day in a randomized manner, such that acclimation to the stress schedule is avoided. This is an important detail as controllability of stressors has been shown to have an impact on the deleterious effects of stress (Christianson et al., 2014). The stressors included alternating periods of food or water deprivation (overnight), continuous cage shaking (1 h on an automatic shaker), social isolation (24 h), forced swim stress (15 min, in 18°C water; limited to three exposures), continuous overnight illumination (12 h), overnight cage-flooding (12 h), exposure to cold temperature (1 h at 4°C), and acute restraint stress (40 min) using plastic DecapiCones (restraint bags) (Iñiguez et al., 2010; Overstreet, 2011).

### 2.3 Western Blot

Protein from NAc tissue punches was isolated using an Illustra TriplePrep kit (GE Healthcare) according to the manufacturer’s instructions and stored at -80°C until use. Protein (10 ug) from each sample was treated with ß-mercaptoethanol and subsequently electrophoresed on precast 4%–10% gradient gels (Biorad), as previously described (Alcantara et al., 2014; Iñiguez et al., 2014). Proteins were then transferred to a polyvinylidene fluoride membrane, washed in 1x Tris-buffered saline with 0.1% Tween-20 (TBST), and blocked in milk dissolved in TBST (5% weight/volume) for 1 h at 25°C. All antibodies were obtained from Cell Signaling (Beverly, Massachusetts) and were used according to the manufacturer’s instructions in 5% bovine serum albumin dissolved in TBST. Blots were probed overnight in 4°C with antibodies against the phosphorylated forms of ERK2, ERK1, MSK, p90-RSK and GAPDH. Separate membranes were probed with antibodies against the total forms of ERK2, ERK1, MSK, p90-RSK and GAPDH was used as a normalizing protein. Membranes were washed several times with TBST and then incubated with peroxidase-labeled goat anti-rabbit IgG (1:10,000; Cell Signaling-Beverly, Massachusetts). Bands were visualized with SuperSignal West Dura substrate (Pierce Biotechnology, Rockford, Illinois), and quantified using ImageJ (NIH). All values are normalized to their respective GAPDH value.

### 2.4 Viral Transfection Surgery, Histology, and Transgene detection

Adult male Sprague Dawley rats were anesthetized with an intramuscular injection of a ketamine/xylazine cocktail (100/10 mg/kg) and given atropine (0.25 mg/kg) subcutaneously to minimize bronchial secretions. Transfection was done via bilateral microinjections (1.0 μl per side over 10 min) of an HSV vector encoding green fluorescent protein (HSV-GFP), HSV-wtERK2, or HSV-dnERK2, in established NAc coordinates (AP: +1.7, Lat: +2.5, DV: −6.7 mm from bregma), angled at 10° from the midline. The construction of the HSV vectors has been previously described (Neve et al., 1997), and the ERK2 virus has been validated both *in vivo* and *in vitro* (Iñiguez et al., 2010; Iñiguez et al., 2019; Warren et al., 2013). Behavioral testing began three days after the viral surgery, a time at which maximal transgene expression has been observed (Iñiguez et al., 2019). No detectable HSV expression was seen in either efferent (e.g., PFC) or afferent (e.g., VTA) regions of the injected area nor in adjacent brain regions (e.g., substantia nigra). Histological assessment of microinjections and transgene detection was performed as previously described (Iñiguez et al., 2019). Briefly, at the end of behavioral testing rats were given an overdose of pentobarbital and perfused transcardially with 0.9% saline, and then cold paraformaldehyde (4%). The brains were removed, post-fixed overnight in 4% paraformaldehyde, and stored in 20% glycerol. Coronal sections (45 μm) were cut on a vibratome and stored in 0.1 M sodium phosphate buffer with 0.05% sodium azide. Slides were then visualized and photographed using a fluorescence microscope and a digital camera. Data obtained from rats with placements outside the intended brain regions (<10% of all experimental animals) were not included in the analyses. A representative hit and infusion image can be seen in Figure 2.

**Figure 1.**
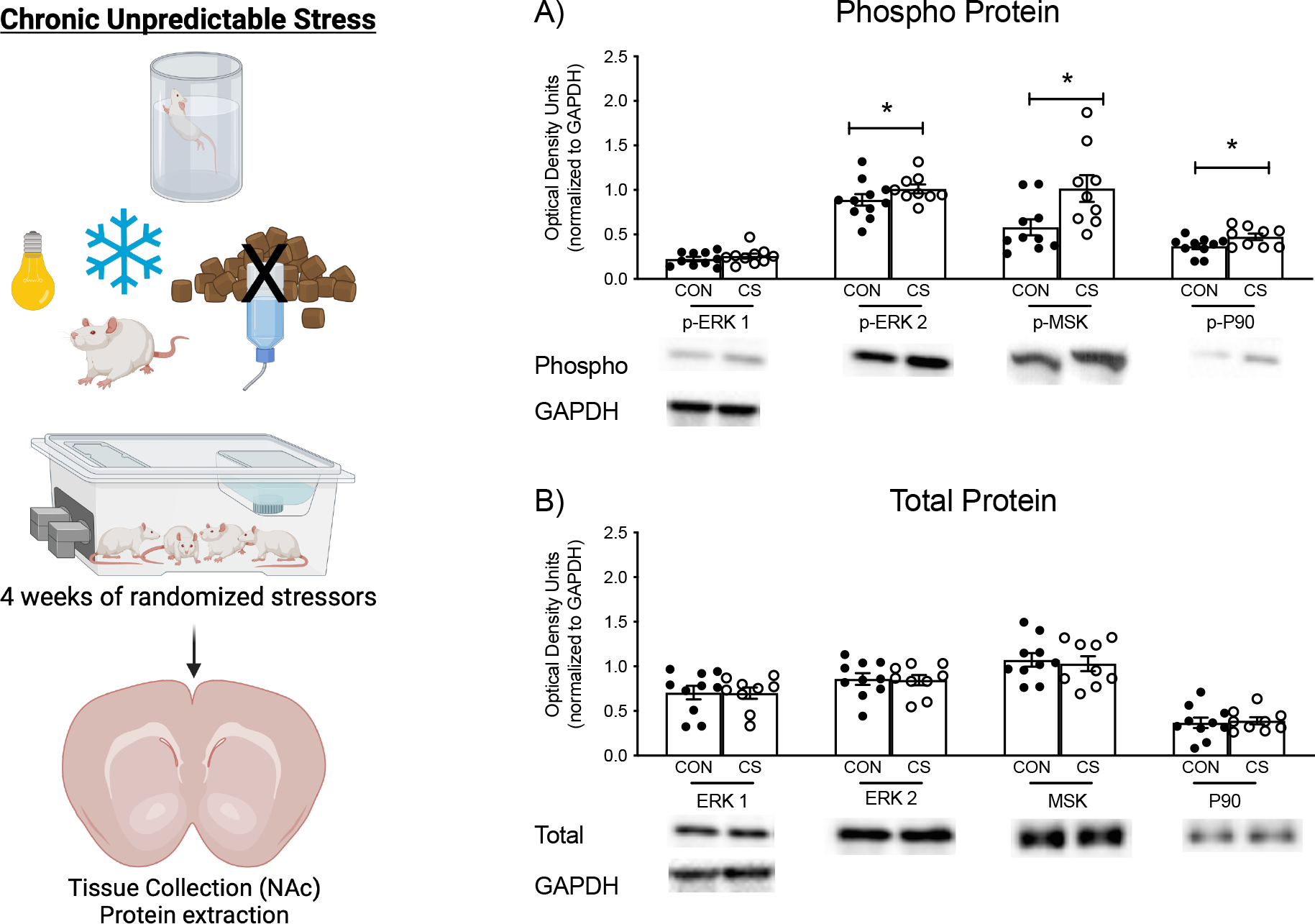
Stress exposure increases phosphorylation of ERK2-related proteins in the nucleus accumbens (NAc). Rats were exposed to four weeks of randomized stressors including overcrowding, alterations to sleep patterns (light cycle, cage tilting, overcrowding), cold exposure, and water or food deprivation. Protein from NAc tissue punches was isolated to assess levels of phosphorylated ERK1, ERK2, MSK, p90-RSK, and GAPDH (A). CS-exposed rats exhibited significantly higher phosphorylated ERK2, MSK, and p90-RSK levels compared to non-stressed controls (CON), while total protein levels remained unchanged (B). Values are normalized to their respective GAPDH values. CS, chronic stress; CON, non-stressed controls; *Significantly different from CON; *p*<0.05.

**Figure 2.**
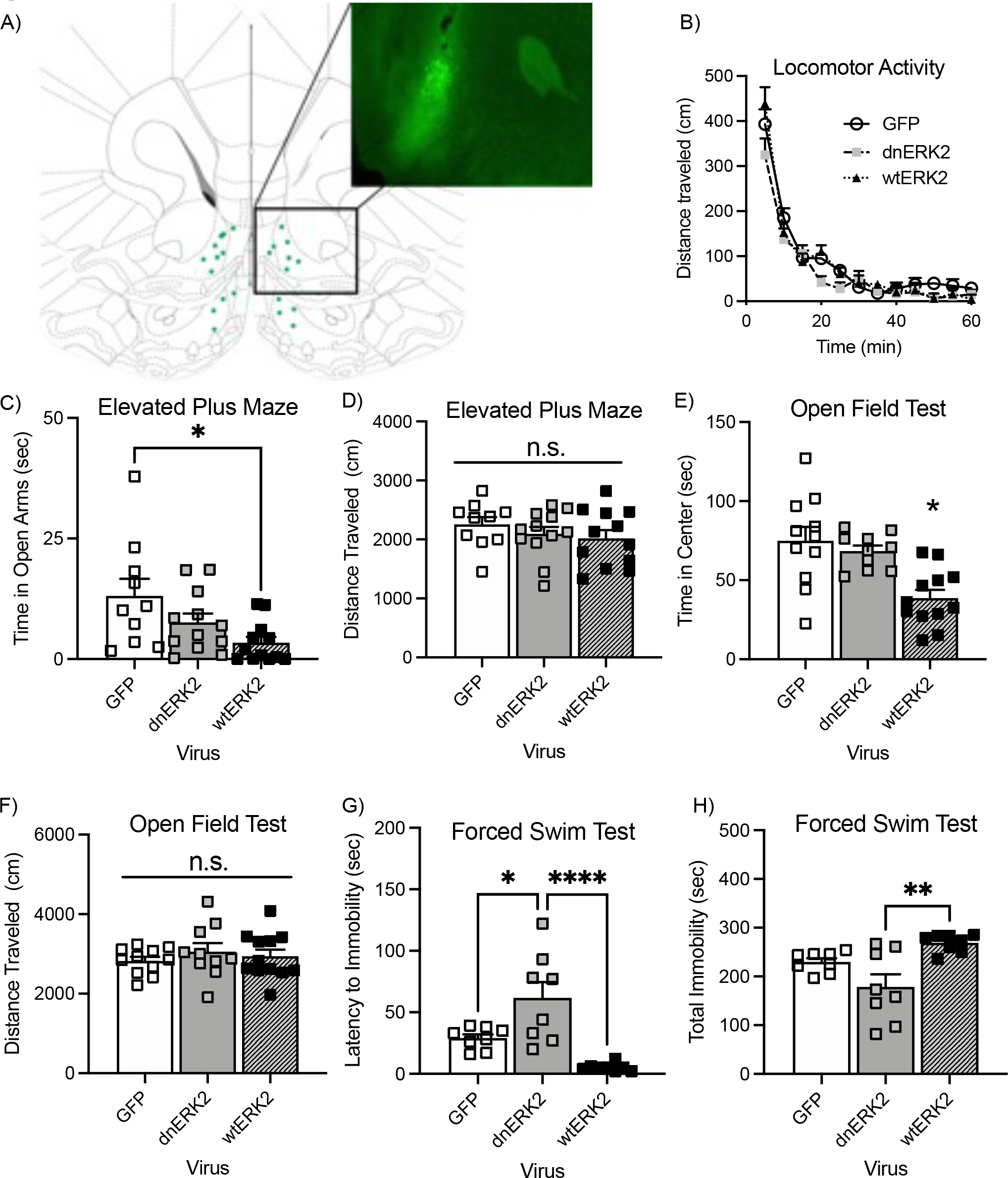
ERK2 modulation in the nucleus accumbens (NAc) regulates anxiety- and depression-like behavior. (A) Schematic of surgical placements within the NAc. (B) No differences were found in total locomotion in rats infused with either HSV-dnERK2 or HSV-wtERK2 when tested in an open chamber locomotion test. However, in the elevated plus maze, HSV-wtERK2 significantly decreased time spent in the open arms (C) without influencing total distance travelled (D). In the open field test, HSV-wtERK2 significantly decreased time spent in the center (E), without influencing total distance travelled (F). Interestingly, on day 2 of the FST, rats infused with HSV-dnERK2 showed a significantly longer latency to immobility compared to both GFP or wtERK2- infused rats (G), with HSV-wtERK2 infusion promoting a corresponding increase in total immobility (H). Interestingly, significantly increased total immobility. \**p*<0.05; error bars indicate ± SEM.

### 2.5 Forced Swim Test

The forced-swim test (FST) is a two-day procedure in which rats are forced to swim under inescapable conditions (Porsolt et al., 1977). On day 1, rats were forced to swim for 15 min. Initially, they engage in escape-like behaviors but eventually adopt a posture of immobility in which they make only the movements necessary to maintain their head above water. When re-tested 24 h later, rats become immobile very quickly; however, antidepressant treatment between the forced-swim exposures can significantly increase escape-like behaviors, an effect that has been used to determine antidepressant efficacy (Lucki, 1997). Rats were initially placed in plastic cylinders (75 × 30 cm) which were filled with 25°C water, as previously described (Iñiguez et al., 2008). Rats were then removed from the water, dried, and placed in a warm enclosure for 30 min. All cylinders were emptied and rinsed between trials. On day two (24 h later), rats were re-tested for five min under identical conditions. Here, the latency to become immobile and total immobility were measured. Latency to immobility was defined as the time at which the rat first initiated a stationary posture and did not attempt to escape from the water (Lucki, 1997). To qualify as immobility, this posture had to be clearly visible and maintained for ≥ 2.0 s (Carlezon et al., 2003).

### 2.6 Basal Locomotor Activity After Day 1 of Forced Swimming

Spontaneous locomotor activity was indexed as distance traveled (in centimeters) in an automated open field box (63 × 63 cm) (Tru Scan, Coulbourn Instruments) in which rats were allowed to freely explore (Iñiguez et al., 2010; Wiley et al., 2009). This behavioral task was performed 24 h after the FST on day 1, to assess whether the effects observed in the FST (day 2) could be confounded by changes in general locomotor activity after viral surgery.

### 2.7 Elevated Plus Maze

The elevated plus maze (EPM) is a behavioral task designed to assess anxiety-like behavior (Hogg, 1996). The maze consisted of two perpendicular, intersecting runways (12 cm wide × 100 cm long) made from gray plastic. One runway had tall walls (40 cm high), termed “closed arms,” while the other had no walls, termed “open arms.” The arms were connected by a central area, and the maze was elevated 1 m from the floor. Testing was conducted between 9:00 A.M. and 1:00 P.M. under controlled light conditions (90 lux), as previously described (Iñiguez et al., 2008; Parise et al., 2020). At the start of the test, rats were positioned in the central area, facing one of the open arms and allowed to explore freely for 5 min, the total time that the rat’s center point (as determined by Noldus Ethovision XT) spent in the open arms along with the total distance travelled, were measured.

### 2.8 Open Field Test

As a second measure of anxiety-like behavior, a separate group of rats were placed in an open field test (OFT) (as described above) for one hour and the time spent in the center of the apparatus (a 31 cm X 31cm square) and total distance travelled were measured during a 60 min period. Time spent exploring the periphery of the arena, versus the center, is a measure of anxiety-like behavior.

### 2.9 Conditioned Place Preference

Conditioned place preference (CPP) was carried out in a three-chamber apparatus as previously described (Alcantara et al., 2014). On the preconditioning day (day 0), rats were allowed to freely explore the entire testing apparatus for 30 min to obtain baseline preference to any of the three compartments (side compartments: 35 cm × 27 cm × 25 cm; middle compartment: 10 cm × 27 cm × 25 cm). Only rats showing no spontaneous preference to either compartment were used (unbiased procedure); this accounted for more than 90% of all the rats tested. Rats then received bilateral microinjections of HSV vectors into the NAc and were allowed to recover for 2 days. After recovery, conditioning trials (2 per day) were given on 2 consecutive days (days 3 and 4). During the conditioning trials, rats received an intraperitoneal (i.p.) saline injection (1 ml/kg) and were confined to one of the compartments of the apparatus for 30 min. After 3 h, rats received cocaine (0, 2.5, 5, or 10 mg/kg, i.p.; Sigma–Aldrich, St. Louis, MO) and were confined to the opposite side compartment for 1 h. Conditioning trials were counterbalanced such that half of the rats received drug in one compartment and the other half received the drug in the opposite compartment. On the final day (day 5), rats were again allowed to freely explore the entire apparatus for 30 min.

### 2.10 Intravenous Catheterization and Intracranial Cannulation

Surgical implantation of jugular catheters and intracranial cannulae occurred in a single surgical procedure under halothane anesthesia (1–2.5%). Silastic catheters pretreated with tridodecylmethyl ammonium chloride heparin were inserted and secured at the jugular vein using Mersilene surgical mesh. Immediately following the catheter implantation, each rat was then placed into a stereotaxic instrument and bilateral guide cannula were inserted into the NAc (AP: +1.7, ML: +/-1.5, DV: -5.7; from bregma). Once inserted, the guide cannula was fixed in place with dental cement and dummy stylets were placed into the guide cannula. Catheters were flushed daily with 0.1 mL heparinized saline and rats were allowed 4–7 days of recovery in their home cage, before experimental procedures began.

### 2.11 Cocaine Self-administration, Extinction and Reinstatement Procedures

Self-administration procedures were performed in operant conditioning chambers (Med-Associates, St. Albans, VT) equipped with two response levers and an infusion pump system. Rats were initially trained to lever press for sucrose pellets on a fixed ratio 1 (FR1) schedule, to facilitate acquisition of cocaine self-administration. After lever-press training, animals were fed ad libitum for at least one day prior to surgery (see above). After recovery from surgery, animals were allowed to self-administer intravenous cocaine (0.5 mg/kg/100 μL injection) on a FR1 reinforcement schedule in daily 2 h sessions for at least three days prior to advancing to an FR5 schedule for an additional five days. Cocaine injections were delivered over 5 s, concurrent with the illumination of a cue light above the active lever and was followed by a 15 s time out when the house light remained off and responding produced no consequence. Inactive lever responses produced no consequence throughout testing. Following self-administration, animals returned to the operant conditioning chambers for daily 6 h extinction sessions. During extinction, responses on the lever previously paired with cocaine (i.e., drug-paired lever) and on the inactive lever were recorded but had no programmed drug or cue delivery. We tested the ability of HSV-wtERK2, HSV-dnERK2 or HSV-GFP control vectors to influence reinstatement of cocaine seeking using a repeated testing design. The day prior to reinstatement testing, rats received bilateral (2.0 μl/side) microinfusions of the vectors into the nucleus accumbens over 10 min through a 33-gauge injection cannulae extending 1 mm beyond the guide cannula. The injectors were left in place for an additional 2 min to allow diffusion into the tissue. Reinstatement testing began 24 h after the infusion and continued for a maximum of four days during over-expression. Each reinstatement session was initiated with 2 h of extinction conditions followed by a 2 h reinstatement test period. Cue-induced reinstatement was measured in the first reinstatement test by the non-contingent (priming) presentation of cocaine-associated cues (2.5 s illumination of cue light and infusion pump, 15 s termination of the house light) delivered every 2 min for the first 10 min. Responding to the drug-paired lever resulted in response-contingent cue delivery throughout the 2 h test (2.5 s illumination of cue light and infusion pump, 15 s termination of house light). Cocaine-primed reinstatement was tested on the next three days by experimenter-administered vehicle or cocaine (5 and 15 mg/kg, intraperitoneal injection) immediately prior to the 2 h reinstatement test in a counterbalanced order. Responses at both the drug-paired and inactive levers were recorded but produced no cue or drug delivery during testing.

### 2.12 Spine Density Analysis

For the viral-based spine analysis, mice were perfused with 4% paraformaldehyde three days after viral microinjection of HSV-GFP, HSV-wtERK2, or HSV-dnERK2 and brains were sectioned at 100 um using a vibratome (Leica VT 1000s). Slices were incubated with and anti-GFP antibody (Millipore), mounted, coded (such that the rater was blind to the experimental conditions) and imaged on a confocal microscope (Zeiss LSM 710). Secondary and tertiary dendrites of NAc MSNs were imaged under a 100× oil-immersion objective at a resolution of 0.027 um × 0.027 um × 0.3 um. Approximately seven to ten neurons were imaged per animal (average total dendritic length = 400 um per animal, *n* = 5–6). Dendritic length was measured using ImageJ (US National Institutes of Health) and spine numbers and types were counted using NeuronStudio (Mount Sinai School of Medicine).

### 2.13 Statistical Analyses

Data analysis was performed using a mixed-design (between and within variables) analysis of variance (ANOVA) followed by Fischer’s *post hoc* tests. When appropriate, Student’s *t*-tests were used to determine statistical significance of planned comparisons. Data are expressed as the mean ± SEM. Reinstatement data were analyzed by mixed design 2-factor ANOVA with repeated measures on extinction baseline vs. cue presentation (Cue reinstatement) or cocaine dose (Cocaine reinstatement). Interaction effects were followed by simple main effects analyses and post hoc tests (Bonferroni’s). Statistical significance was defined as *p*<0.05.

## 3. Results

### 3.1 Western Blot

Protein was extracted from NAc tissue punches to assess stress-induced changes in ERK2 and its related signaling molecules. A Student’s *t*-test revealed differences in phosphoprotein expression between CUS-exposed rats and their CON counterparts (n=9–10/group; Figure 1A). Specifically, increased phosphorylated levels of ERK2 (t_17_=2.246, *p*=0.0382), MSK (t_17_=2.557 *p*=0.0204), and p90RSK (t_17_=2.193, *p*=0.0425) were observed in the NAc as a function of stress exposure. Levels of phospho-ERK1 (t_17_=.2804, *p*=0.7826) were not significantly different from non-stressed controls. Separate membranes were probed for the total forms of the proteins of interest and no significant differences were observed between stress and non-stressed controls (n= 9–10/group; Figure 1B) [ERK2 (t_17_=0.1422, *p*=0.8886); MSK (t_17_=0.3765 *p*=0.7112); p90RSK (t_17_=0.3004, *p*=0.7675); ERK1 (t_17_=.06039, *p*=0.9526)].

### 3.2 Elevated Plus Maze

To determine the role of ERK2 signaling in the NAc on anxiety-like behavior, a group of rats were microinjected with HSV-GFP, HSV-dnERK2, or HSV-wtERK2 and three days later exposed to the EPM. Time spent in the open arms of the EPM (*F*_(2, 31)_=4.5; *p*<0.05, Figure 2C) varied as a function of viral pre-exposure. More specifically, HSV-dnERK2 significantly decreased time spent in the open arms compared to HSV-GFP rats (*p*<0.05). Neither HSV-dnERK2, nor HSV-wtERK2 influenced total distance travelled in the EPM compared to HSV-GFP rats (Figure 2D).

### 3.3 Open Field Test

To further characterize the effect of stress exposure on anxiety-like behavior, a separate group of rats was exposed to the OFT after receiving microinjections of HSV-GFP, HSV-dnERK2, or HSV-wtERK2 into the NAc. Time spent in the center of the OFT varied as a function of viral pretreatment (*F*_(2, 30)_=9.9; *p*<0.01; Figure 2E). HSV-dnERK2, but not HSV-wtERK2, significantly decreased the time spent exploring the center of the OFT arena compared to HSV-GFP rats. To test for changes in locomotion as a result of stress exposure, total distance travelled during the OFT was measured. No differences in total locomotion were seen in the HSV-dnERK2 or HSV-wtERK2 exposed rats compared to HSV-GFP rats (*p*>0.05; Figure 2F), suggesting that locomotor activity is not influenced by viral attenuation or overexpression.

### 3.4 Forced Swim Test

To assess for stress sensitivity, HSV-GFP, HSV-dnERK2, or HSV-wtERK2 were microinjected into the NAc of rats and three days later, their behavior in the FST was analyzed (Figure 2). Latency to immobility varied as a function of viral pretreatment (*F*_(2, 21)_=13.8, *p*<0.01; Figure 2G). Total immobility time also varied as a function of viral pretreatment (*F*_(2, 21)_=8.2, *p*<0.01; Figure 2H). Post hoc analyses revealed that HSV-dnERK2 significantly increased latency to become immobile and significantly decreased total immobility, while HSV-wtERK2 significantly decreased latency to become immobile compared to HSV-GFP exposed rats. Furthermore, HSV-wtERK2 significantly reduced latency to become immobile and significantly increased total immobility compared to HSV-dnERK2 rats.

### 3.5 Basal Locomotor Activity after Day 1 of Forced Swimming

To test for changes in locomotion, a separate group of rats was injected with HSV-GFP, HSV-dnERK2, or HSV-wtERK2 prior to day 1 of the FST. Twenty-four hours later, these rats were placed into the open field and total distance travelled was measured (Figure 2B). No differences in locomotion were found among any of the groups (*p*>0.05).

### 3.6 Conditioned Place Preference

The effects of HSV-GFP, HSV-dnERK2dn, and HSV-wtERK2treatments on cocaine (0, 5, and 10 mg/kg) CPP are shown in (Figure 3A). Time spent in the cocaine-paired compartment varied as a function of virus-(*F*_(2,57)_=4.2, *p*<0.05) and drug-treatment (*F*_(2,57)_=7.3, *p*<0.01), as well as their interaction (virus × drug interaction: *F*_(6,85)_=2.7, *p* <0.05). Rats that received HSV-GFP, HSV-wtERK2, or dnERK2 and were conditioned to saline did not show preferences for either side of the compartments. Conversely, rats receiving HSV-wtERK2 microinjections into the NAc spent significantly more time in environments paired with moderate doses (2.5 and 5 mg/kg) of cocaine (*p*<0.05), whereas rats receiving HSV-dnERK2 microinjections did not consistently approach the cocaine-paired environments (*p*>0.05).

**Figure 3.**
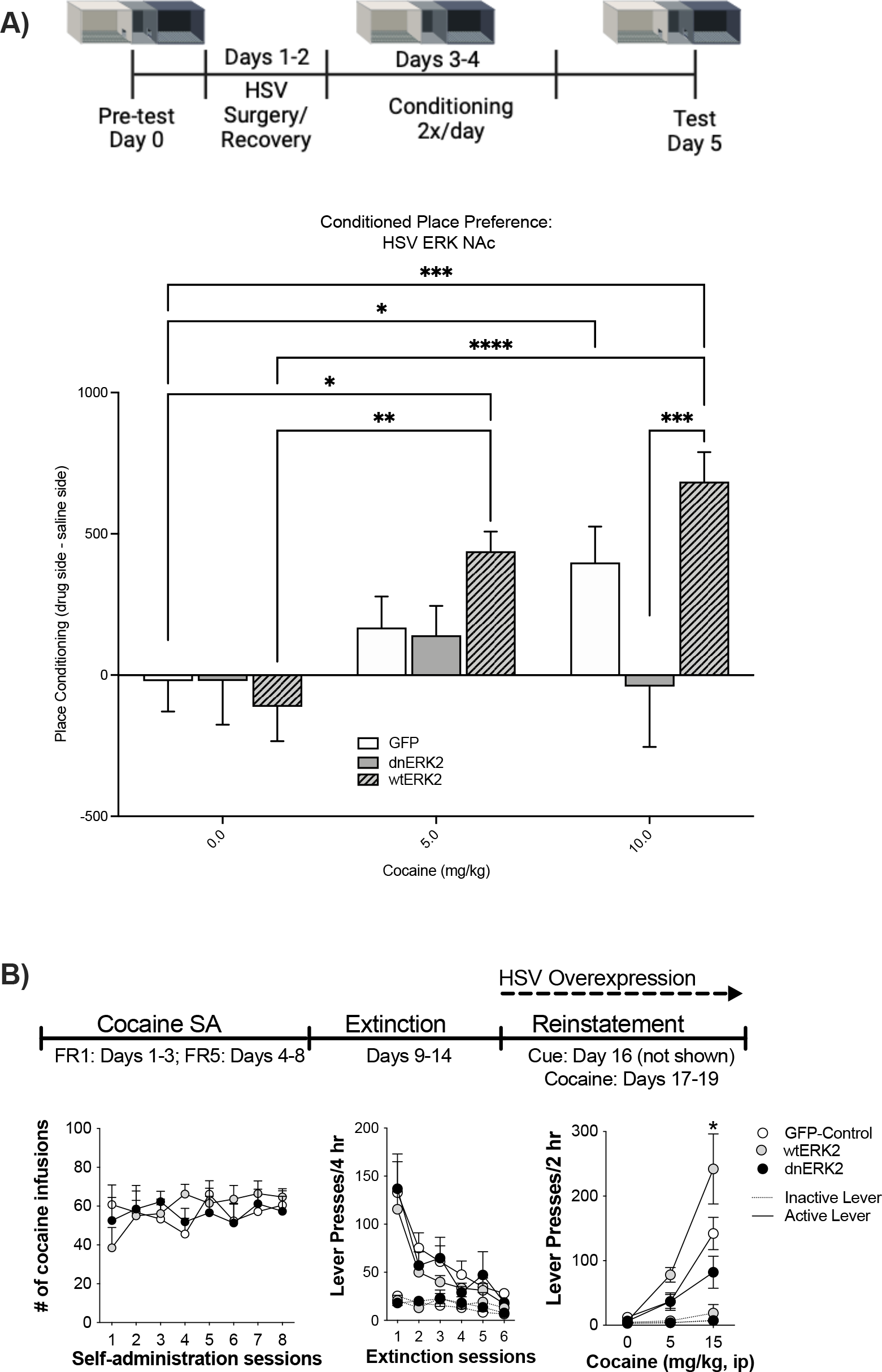
ERK2 modulation in the nucleus accumbens (NAc) regulates cocaine reward. (A) Cocaine-mediated place conditioning protocol. HSV-GFP rats showed reliable CPP to 10.0 mg/kg of cocaine but not 5.0 mg/kg. Interestingly, HSV-wtERK2 rats showed robust place preference at both 5.0 and 10.0 mg/kg of cocaine, whereas HSV-dnERK2 blunted CPP acquisition to both doses. (B) Experimental timeline for self-administration, extinction training, and reinstatement testing during HSV over-expression. Over-expression of dnERK2 or wtERK2 produced bidirectional effects on cocaine seeking induced by cocaine priming. \**p*<0.05; error bars indicate ± SEM.

### 3.7 Cocaine Self-administration, Extinction, and Reinstatement

Cocaine self-administration and extinction responding were balanced across treatment groups, and animals were infused with either HSV-GFP, HSV-wtERK2, or HSV-dnERK2 into the NAc 1 day prior to the onset of reinstatement testing (Figure 3B). Neither wtERK2 nor dnERK2 directly altered cue-induced reinstatement of cocaine seeking (Group x Cue: *F*_(2,26)_; *p*>1). However, wtERK2 markedly enhanced, while dnERK2 attenuated, cocaine-induced seeking compared to GFP-expressing controls (Group x Cocaine Dose: *F*_(4,52)_=3.72, *p*<0.01). These findings indicate that enhancing ERK2 function in the NAc may exacerbate the risk of cocaine relapse.

### 3.8 Spine Density

To determine whether HSV-GFP, HSV-wtERK2, or HSV-dnERK2 injections into the NAc would alter synaptic plasticity, we measured spine density within the NAc. Spines were categorized based on their distinct phenotypes and interestingly, we observed that mushroom (*F*_(2, 16)_=3.93; *p*<0.05, Figure 4A) and stubby (*F*_(2, 16)_=3.6; *p*<0.05, Figure 4C), but not thin (*F*_(2, 16)_=0.2663; *p*<0.05, Figure 4B) spines were significantly influenced by viral pretreatment. More specifically, HSV-wtERK2-infused rats had a significantly increased density of both mushroom and stubby spines compared to HSV-GFP-infused rats *(p*<0.05).

**Figure 4.**
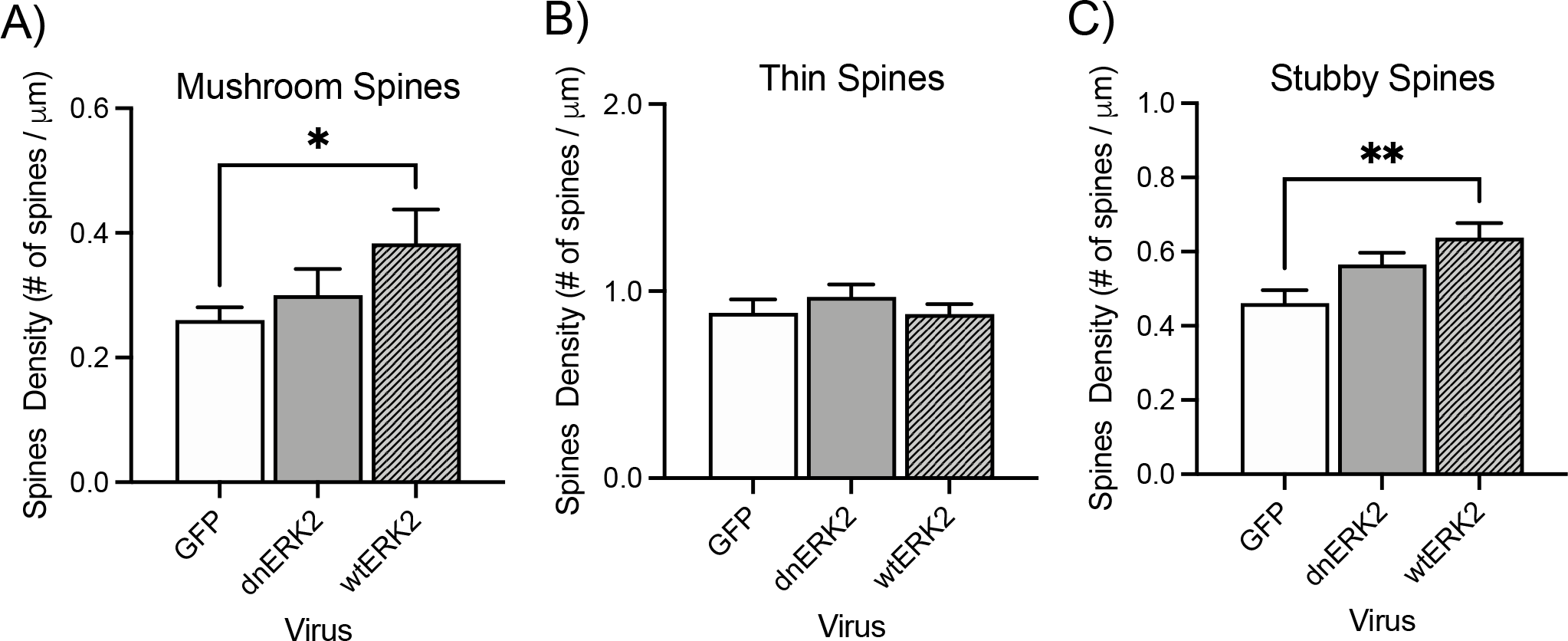
ERK2 activity in the NAc regulates synaptic plasticity. HSV-wtERK2 significantly increases mushroom (A) and stubby (C) spine density without influencing thin spines (B). *Significantly different from HSV-GFP;\**p*<0.05; error bars indicate ± SEM.

## 4. Discussion

The present study was designed to investigate the role of ERK2 in stress responses and to assess whether virally-mediated over-expression or knockdown of ERK2 within the NAc is sufficient to influence behavioral assessments of mood or drug reward. Here, we show that modulation of ERK2 in the NAc influences both behavioral measures of anxiety- and depression-like behavior as well as sensitivity to the reinforcing properties of cocaine. Specifically, our findings demonstrate that adult rats exposed to four weeks of CUS exhibit an increase in ERK2 protein expression accompanied by corresponding increases in its related signaling molecules. Using a viral-mediated approach to mimic stress-induced expression, we show that increasing ERK2 activity in the NAc decreased latency to immobility in the FST while also decreasing overall effort in the task (i.e., increased total immobility). Additionally, we observed an anxiogenic response to the EPM exemplified by a decrease in the time spent in the open arms of the EPM. Similarly, we observed less time spent in the center of the OFT following overexpression of ERK2 in the NAc. Together, these deficits are indicative of increased susceptibility to stress-eliciting stimuli as a direct function of increased levels of the ERK2 protein. Interestingly, decreasing ERK2 signaling within the NAc increased latency to immobility and decreased total time immobile in the FST without producing a significant impact on anxiety-like behaviors in the EPM or OFT. These results are not necessarily contradictory. While the FST has been traditionally used to identify potential antidepressant efficacy of pharmacological or behavioral manipulations, other tests tend to be more variable in assessing affective behavior. Additionally, antidepressant drugs can reduce total immobility in the FST, while increasing anxiety-like behavior in other measures (Iñiguez et al., 2010; Oh et al., 2009). Indeed, it may take up to one month of continuous antidepressant treatment for anxiolytic effects to be seen (Iñiguez et al., 2010; Oh et al., 2009), thus, given the short-term testing window of our experimental condition, it is not unreasonable to see a range of anxiety-related behaviors.

Chronic and acute exposure to cocaine have been shown to alter neurotrophin signaling within the NAc, initiating activity of downstream effectors, including ERK2 (Berhow et al., 1996; Bolaños and Nestler, 2004). We now expand on this work and highlight that directly increasing ERK2 expression not only promoted stress-susceptibility but also shifted the rewarding threshold of cocaine (Iñiguez et al., 2010; Iñiguez et al., 2010; Warren et al., 2011). Conversely, we show that decreasing ERK2 activity reduced depression-like behavior and blunted sensitivity to cocaine reward (Iñiguez et al., 2010; Iñiguez et al., 2010).To better understand the motivating and reinforcing effects of ERK2 modulation in the NAc, rats were trained to self-administer cocaine and then placed through extinction training prior to viral transfection and reinstatement testing. Reinstatement of cocaine seeking was found to be strongest in HSV-wtERK2-infused rats. This is particularly interesting since previous studies have shown that stress exposure can influence both the acquisition and extinction of drug-associated behaviors. Particularly, exposure to acute and chronic stress has been shown to re-establish methamphetamine or cocaine seeking in rats, even after an extended period of extinction (Manvich et al., 2016; Taslimi et al., 2018). Furthermore, studies have shown that stress, experienced at various stages of life, can influence the acquisition of cocaine self-administration and impede the extinction of cocaine-seeking behaviors (Shinohara et al., 2018; Wong and Marinelli, 2015). Considering that both the acquisition and extinction of these behaviors are extended learning processes spanning many days, the short temporal dynamics of the HSV constructs make investigating ERK2’s role in facilitating either of these behaviors difficult. Therefore, future studies utilizing longer lasting viral manipulations, such as adeno-associated viruses, are critical to understanding the role of ERK2 in long-term drug-associated behaviors. Nevertheless, previous work does support the notion that ERK2 can contribute to these drug processes. For example, activation of a particular subpopulation of heteromeric D1-D2 MSNs in the NAc has been shown to attenuate ERK2 signaling and consequently abolish cocaine CPP by affecting both acquisition and extinction of cocaine seeking (Hasbi et al., 2017). Furthermore, the distribution of spine type and complexity within the NAc have also been found to be influenced by cocaine and stress (Christoffel et al., 2010; Christoffel et al., 2011; Dietz et al., 2012; Warren et al., 2014). Indeed, given the established role of ERK2 in memory and synaptic plasticity (Adams and Sweatt, 2002; Miller and Marshall, 2005; Radulovic, and Tronson, 2010), we were also interested in determining whether direct manipulation of ERK2 could alter dendritic spine density thereby, mediating some of the observed behaviors. To this end, we examined alterations in dendritic spine density among MSNs after viral overexpression of either HSV-wtERK2 or -dnERK2 in the NAc. Our findings revealed that only HSV-wtERK2 led to a significant increase in dendritic spine density compared to HSV-GFP-exposed rats. Further assessment revealed an enhancement of more mature spine types with preferential enrichment of stubby and mushroom spines versus thin spines. Interestingly, the upregulation of ERK2 elicited greater deficits in tasks that rely on learning and memory; arguably. Notably, CPP, the reinstatement of drug seeking, and to a lesser extent, behavior in the FST, all rely on learning and memory processes (Huston et al., 2013; Sun and Alkon, 2004; West, 1990). In contrast, the EPM and OFT do not require memory formation and were found to be more strongly influenced by HSV-dnERK2 modulation (Sharma and Kulkarni, 1992). It is possible, then, that the increased ERK2 activity may not necessarily make cocaine more rewarding in wtERK2-exposed rats, but instead, it might enhance the formation of associations with cues and contexts linked to cocaine use. This potential mechanism offers valuable insights into ERK2’s role in the habitual aspects of drug addiction and mood disorders. For example, individual differences in ERK2 signaling could contribute to susceptibility to drug-associated or stress-triggering environments by selectively enhancing reward-related learning processes through the NAc. Nevertheless, subsequent studies are needed to determine the role of ERK2 in NAc-facilitated learning of drug- and stress-related memories. It is worth noting that his study has certain limitations. While HSV-mediated modulation provides an ideal way to investigate the specific role of ERK2, it may not fully recapitulate the broader biological consequences of chronic stress exposure. Additionally, given the temporal restriction of HSV expression, the behavioral testing window in this experimental design is limited.

## Conclusions

Here, we demonstrate a role for neurotrophin-related signaling within the NAc in mediating mood-related and cocaine-seeking behavior. Viral over-expression of ERK2 in the NAc increased depression-related and anxiety-like behavior, altered sensitivity to cocaine relapse, and increased dendritic spine density within the NAc. In contrast, knocking down ERK2 in the NAc decreased depression-like behavior and cocaine sensitivity. Taken together, these data suggest an overlapping role for ERK2 signaling in the NAc for mediating sensitivity to mood-related stimuli and drugs of abuse, underscoring the need for a deeper understanding of second messenger signaling in the NAc, an area aptly suited for the integration of drug addiction and affective dysregulation.

## Author contributions

LFP and SDI conceived the experimental design. LFP, SDI, BLW, and EMP performed the viral infusion surgeries and subsequent behavioral testing. LFP and SDI performed the conditioned place preference tests and RKB and DD conducted the self-administration tasks. EJN and CAB helped with data analysis and interpretation. LFP and SDI wrote the final manuscript.

## Data availability statement

Data available upon reasonable request to the corresponding author.

